# Remodeling and activation mechanisms of outer arm dyneins revealed by cryo-EM

**DOI:** 10.1101/2020.11.30.404319

**Authors:** Shintaroh Kubo, Shun Kai Yang, Corbin Black, Daniel Dai, Melissa Valente, Jacek Gaertig, Muneyoshi Ichikawa, Khanh Huy Bui

**Affiliations:** Department of Anatomy and Cell Biology, McGill University, Montréal, Québec H3A 0C7, Canada; JSPS Overseas Research Fellow; Department of Cellular Biology, University of Georgia, Athens, GA 30602, USA; Department of Systems Biology, Graduate School of Biological Sciences, Nara Institute of Science and Technology, Ikoma, Nara 630-0192, Japan; PRESTO, Japan Science and Technology Agency, Kawaguchi, Saitama, 332-0012, Japan; Centre de Recherche en Biologie Structurale, McGill University, Montréal, Québec H3A 0C7, Canada

**Keywords:** Cilia, Doublet microtubule, Outer arm dynein, Cryo-electron Microscopy

## Abstract

Cilia are thin microtubule-based protrusions of eukaryotic cells. The swimming of ciliated protists and sperm cells is propelled by the beating of cilia. Cilia propagate the flow of mucus in the trachea and protect the human body from viral infections. The main force generators of ciliary beating are the outer dynein arms (ODAs) which attach to the doublet microtubules. The bending of cilia is driven by the ODAs’ conformational changes caused by ATP hydrolysis. Here, we report the native ODA complex structure attaching to the doublet microtubule by cryoelectron microscopy. The structure reveals how the ODA complex is attached to the doublet microtubule via the docking complex in its native state. Combined with molecular dynamics simulations, we present a model of how the attachment of the ODA to the doublet microtubule induces remodeling and activation of the ODA complex.

## Introduction

Motion is an important aspect of life. In eukaryotes, cilia and flagella are responsible for cell motility. These microscopic hair-like organelles bend several tens of times per second to generate fluid flows. Cilia in trachea generate the flow of mucus and protect our body from infectious agents such as viruses. There is a canonical axonemal 9+2 structure where the central pair microtubules are surrounded by nine doublet microtubules (Fig. 1A). The propulsive force generators of cilia and flagella are the axonemal dyneins. As the molecular motors, the axonemal dyneins drive the sliding of doublet microtubules, which is then converted into the bending of the cilium. The axonemal dyneins consist of outer arm dyneins and inner arm dyneins. The outer dynein arm (ODA) regulates the beat frequency while the inner arm dyneins are important for the waveform of the cilium (Brokaw, C. J. and Kamiya, R., 1987). Improper assembly of the ODA complex causes ciliopathies in humans (reviewed in Reiter, J. F. and Leroux, M. R., 2017).

**Fig 1.**
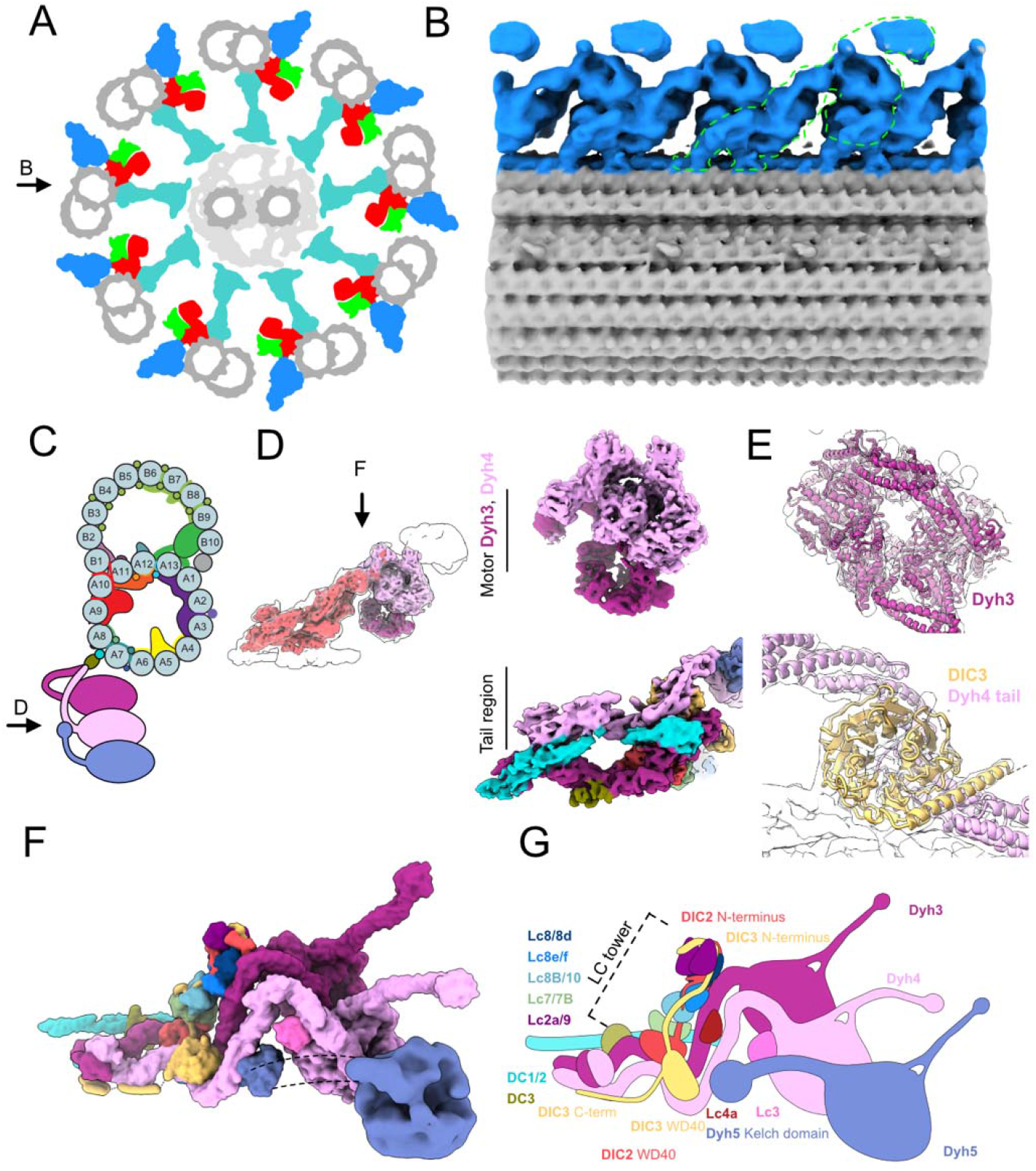
Cryo-EM structure of the ODA complex on the doublet microtubule. (A) Schematic diagram of axoneme structure of cilia viewed from the base of the cilia. Doublet microtubule: gray; outer dynein arm: blue; inner dynein arm: red; dynein regulatory complex: green; radial spokes: cyan. Black arrow indicates the view in B. (B) The 24-nm structure of the doublet microtubule from K40R mutant filtered to 18 Å showing the row of the ODA. The green outline indicates the single ODA complex. (C) A schematic cartoon of the doublet microtubule and the ODA complex. Arrow indicates the view in D. (D) The focused refined maps of the tail and the heads of the ODA are shown within the map of the entire ODA complex (left). The focused refined maps of Dyh3 and Dyh4 heads and the tail (right). (E) Fitting of models into maps (Dyh3, top; DIC3 and Dyh4 tail, bottom). The α-helix part of the DIC3 was more structured in our map compared with that of the Shulin-ODA complex. (F, G) The surface render of the model (F) and the schematic cartoon (G) of the ODA with the docking complex. The Dyh5 is too flexible to resolve well by cryo-EM. The Dyh5 head is only shown as the 18-Å resolution surface render and the Dyh5 tail is drawn as dotted lines. All the components (ICs, LCs and DCs) are colored and indicated and will be used consistently in all the figures. C-terminal side of the DIC3 has more structured region along the Dyh4 HC tail on the doublet.

Unlike cytoplasmic dyneins which walk on microtubules while carrying cargos, axonemal dyneins are anchored firmly on the doublet microtubules (Bui, K. H. et al., 2008; Bui, K. H. et al., 2009). The ODA complex is composed of two or three heavy chains (HCs) depending on the species (Diamant, A. G. and Carter A. P., 2013), two intermediate chains (ICs) and numerous light chains (LCs). For example, human cilia have two headed ODA (DNAH5 and DNAH9) while ODA of Tetrahymena contain three heavy chains (Dyh3, 4 and 5). Dynein HCs are the most important force producing components and each HC has a head domain and a tail domain. The three head domains are coupled together at the level of the tail domain. Within the head domain of each ODA HC, there is an AAA+ ring (composed of AAA1 to AAA6 subdomains) where ATP is hydrolyzed to exert force. The ODA complex is stably attached to the A-tubule of the doublet microtubule and produces force while interacting with the B-tubule of the neighboring doublet microtubule. The docking of the ODA complex to the A-tubule of the doublet is mediated by the HC tail domain. The interaction with B-tubule of the neighboring doublet is mediated by the microtubule binding domain (MTBD) at the tip of the stalk which extends from the AAA+ ring. ICs and LCs have various regulatory roles (reviewed in King, S. M., 2012). Most of the ICs and LCs interact with the tail domain. However, LC1 is bound to the MTBD of the Dyh3 head (Ichikawa, M. et al., 2015; Toda, A. et al., 2020).

By cryo-electron tomography (cryo-ET) work, ODAs were shown to form a 24-nm repeating row on the doublet microtubules (Bui, K. H. et al., 2012; Lin J. and Nicastro, D. 2018). The ODA complex is proposed to be attached via the docking complex (DC) (Owa, M. et al., 2014; Oda, T., et al., 2016). However, the interactions among the ODA and DC complexes and the doublet have not been revealed in subnanometer resolution. Within each ODA complex, the three heads are aligned parallel to each other so that all three head domains can interact with the adjacent doublet microtubule in a proper orientation. The architecture of ODA complex has been studied by negative staining EM using purified ODA complexes (Ichikawa, M. et al., 2015), or cryoelectron tomography (cryo-ET) of the intact axoneme (Oda, T. et al., 2016). However, the resolutions were limited in these studies and the modeling of the protein subunits was not possible. Recently, a cryo-EM structure of the inactive ODA complex (before its incorporation into cilia) was obtained (Mali, G. R. et al., 2020). In the inactive form, three head domains are packed together by a regulator protein, Shulin. This conformation is markedly different from the active conformation observed in the axoneme where the three heads are in parallel arrangement. The subunit architecture was also revealed in high resolution for the inactive Shulin-ODA complex (Mali, G. R. et al., 2020). To understand how ODA is incorporated into the axoneme structure and activated, it was crucial to obtain a high-resolution structure of the ODA complex in the context of the doublet microtubule.

Here, we revealed the ODA complex on the doublet microtubule at 5.5-7 Å resolutions. Our structure showed in detail how the ODA complex is attached to the doublet, including its interaction with the DC complex. Combined with molecular dynamics (MD) simulations, we have revealed how the ODA complex undergoes an activating rearrangement when it is docked onto the doublet microtubule.

## Results

### Cryo-EM structure of the ODA attached to the doublet microtubule

In our previous studies of the doublet microtubule structures, the ODA complexes were removed by high-salt treatment from doublets (Ichikawa, M. et al., 2017; Ichikawa et al., 2019; Khalifa, A.A.Z. et al., 2020). Here, to obtain the native structure of the ODA complex on the doublet, we tried to obtain a wild-type (WT) Tetrahymena doublet microtubule structure without a salt wash (Fig. S1A). The individual doublet microtubules were separated from the rest of the axoneme induction of microtubule sliding using ATP. However, in the disintegrated WT axonemes, the ODAs tended to detach from the doublet (Fig. S1B) and we were not able to obtain the high resolution structure of the ODA by single particle analysis (Fig. S1D and E). We noticed that the sliding of doublets in the ATP-reactivated K40R (Gaertig et al., 1995) or MEC17-KO mutant *Tetrahymena* axonemes (Akella, J. S. et al., 2010) was less efficient as compared to the WT Tetrahymena axonemes, possibly due to different levels of post-translational modifications of tubulin. This enabled us to obtain cryo-EM images of partially split doublet microtubules with attached ODA complexes (Fig. S1C). Using the cryo-EM images, we first obtained a 24-nm doublet microtubule unit structure as previously described (Ichikawa, M. et al., 2017; Ichikawa, M. et al., 2019; Khalifa, A. A. Z. et al., 2020) at a 3.9 Å resolution (Fig. S2A). However, the resolution of the ODA part was not high enough due to its flexibility. Therefore, we performed signal subtraction of the doublet and obtained the ODA structure with a 7.8 Å resolution. From here, different regions of the ODA were further revealed by focused refinement. The tail region’s resolution was improved to 5.5 Å, the Dyh3 head part to 5.8 Å and Dyh4 head part to 7 Å (Fig. S2B). The resolution of the Dyh5 head remained at 17 Å since it was furthest from the doublet and more flexible. The head domains were in a parallel configuration (Fig. 1B) as previously shown by cryo-ET studies (Bui, K. H. et al., 2008; Lin J. and Nicastro, D. 2018). Our structure fits well with previous cryo-ET structure of the intact axoneme, thereby confirming it is a physiological structure of the ODA complex (Fig. S1F). Using the recent Shulin-ODA complex structure from Tetrahymena as a starting model, we were able to build in all the components of the ODA complex except for Dyh5’s head (Fig. 1C-G). Since Dyh5 is not conserved in vertebrates (Ueno, H. et al., 2014; Lin, J. et al., 2014), an atomic model for conserved part of the ODA complex on the doublet was obtained. The ICs and LCs were also assigned to the density map. The majority of LCs were forming the LC tower as earlier observed in the Shulin-ODA complex (discussed below). Compared with the Shulin-ODA complex, there were some parts of ICs which were structured in our map and modeled (Fig. 1E-G). There was a density not observed in the Shulin-ODA complex associated with Dyh4, and LC4 was tentatively assigned to this region (Fig. 1F and G). Very recently, Tetrahymena ODA model based on cryo-EM structure of reconstituted ODA array on the doublet was reported (Rao, Q. et al., 2020). Our model was similar with obtained reconstituted ODA complex validating our model. In our structure, there was also an additional density of two segments of coiled coil with a globular domain at the end which is running along the tail domain of the ODA (cyan parts in Fig. 1D, F and G) (discussed later).

### Attachment of the ODA complex to the doublet via the docking complex

The part of DC on the doublet was also visualized in our structure which allowed us to understand how the ODA complex is attached to the doublet (Fig. 2 and Fig. S3A). We identified Tetrahymena proteins Q22T00, Q233H6 and I7M2C6 as DC1/2 and 3 following the names of the counterparts of Chlamydomonas (Fig.S3B and C, see Materials and Methods for details). Along the doublet, DC1/2 forms a 24-nm repeating unit, the same periodicity as the repeating unit of the ODA complex. DC3 is associated with the DC1/2 coiled coil region on the doublet serving as a marker for the ODA attachment (Fig. 2C and Fig. S3A) as previously proposed (Ma, M. et al., 2019). The α-helix bundle-3 of Dyh3-HC was interacting with the DC3 (Fig. 2C). The tethering of the ODA to the doublet microtubule is mediated by Dyh3 alone and ICs/LCs are not involved with the microtubule. The DC1/2 also appears to connect to the coiled-coil density with a gap associated with the neighboring ODA tail domain (dashed line in Fig. 2A, and Fig. S3H). This model is consistent with the coiled coil prediction of DC1/2 with a gap (Fig. S3B) and the previous tagging results of DC2 of Chlamydomonas reinhardtii by cryo-ET in Oda, T., et al., (2016) (Fig. S3D-G). Therefore, the density associated with the tail domain corresponds to the C-terminal side of DC1/2 (Fig. S3K).

**Fig 2.**
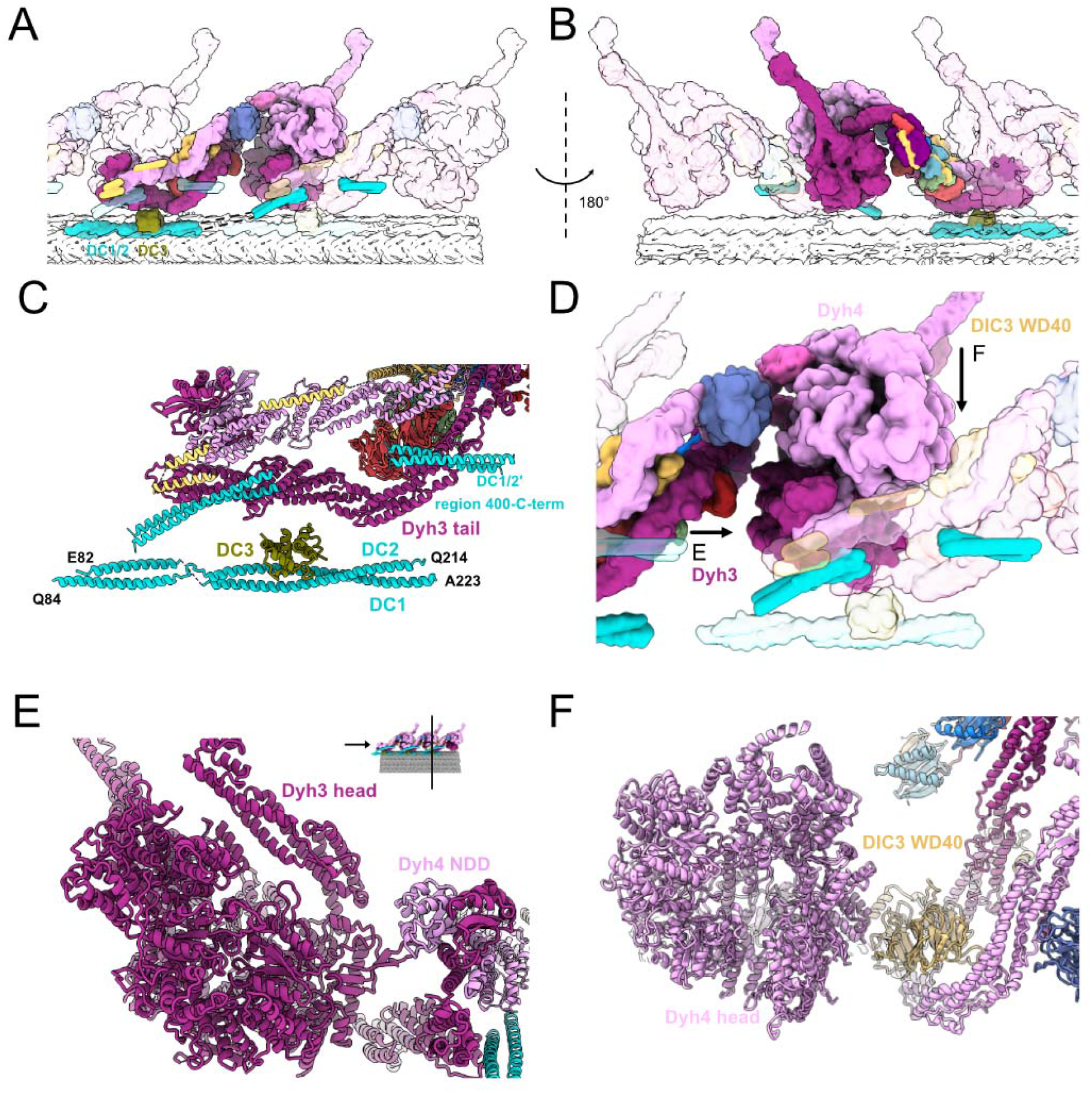
Interactions between the ODA complex and the DC. (A, B) The stacking of the ODA complex in the axoneme viewed from inside (A) and outside (B) the cilia. All the components belonging to the middle ODA complex are shown in color. The proximal and distal ODA complexes are shown in transparent. (C) Model of the DC and the tail of the ODA. (D) The interaction of the head of the ODA complex with the tail of the next ODA complex unit. Dyh3 head interacts with the dimerization domain NDD of the proximal Dyh4. The Dyh4 head interacts with the DIC3 WD40 of the proximal ODA complex. Arrows indicate the views in (E) and (F). (E) Model view of the interaction between the Dyh3 head with the dimerization domain NDD of the proximal ODA complex. (F) Model view of the interaction between the Dyh4 head with the DIC3 WD40 of the proximal ODA complex.

The ODA complex is anchored to the doublet by the above-mentioned two sites: DC1/2 and DC3. Over these docking sites, there is a layer composed of Dyh3 HC, DIC2’s WD domain, LC tower and the C-terminal side of DC1/2. Another layer of Dyh4 and DIC3 is further built onto this layer (Fig. 2A and B). The three head domains are stacked together in a parallel configuration. The stacking of both the tail and the head domains is important for the proper attachment of the ODA onto the doublet.

While the ODA complex is not directly in contact with the doublet microtubule in its native state, ODAs can be reconstituted on preassembled singlet microtubules or doublets lacking DCs with 24-nm periodicity (Ueno, H. et al., 2008; Owa, M. et al., 2014). Our structure suggests that the ODA can be reconstituted on the microtubule without DCs by the interaction of the ODA heads with the proximal neighboring ODA tail domains (Fig. 1B and Fig. 2D-F). More specifically, the head of Dyh3 seems to interact with the N-terminal dimerization domain (NDD) of the proximal ODA Dyh4 tail while the Dyh4 head interacts with DIC3 WD40 domain of the proximal ODA. These interactions probably enable the spontaneous alignment and stacking of the ODA complexes on the doublet microtubule without the help of the DCs. Similar tail-to-head interaction was recently observed in reconstituted ODA array on the doublet microtubule (Rao, Q. et al., 2020). Since our structure is the native structure of ODA on the doublet, our structure validated their model of the interaction.

### Remodeling of the tail and the head

It has been recently proposed that the ODA complex, before its assembly on the doublet, is in an inactive form with the regulatory protein Shulin (Mali, G. R. et al., 2020). To understand how ODA rearranges into an active form on the doublet, we compared the structures of the Shulin-ODA complex with our ODA complex on the doublet (Fig. 3 and Fig. S5). The overall appearance of the active ODA complex on the doublet was quite different from that of the inactive Shulin-ODA complex, which has a more elongated conformation (Fig. 3A). The elongated conformation of the inactive Shulin-ODA complex stems from the positions of the motor domains which are fixed in pre-powerstroke conformations (discussed later). The most unchanged region was the base region of the Dyh4 HC up to the bundle-4 region where the DIC3’s WD40 domain resides (Fig. 3B and Fig. S5A). Dyh3 HC, DIC2 and LC tower were rotating together about ∼90 degree relative to the Dyh4’s base region (Fig. 3B). With this configuration change, Dyh3 and Dyh4 HCs are no longer crossed to each other and freed. In the active conformation, DIC2’s WD40 domain interacts with Dyh4 HC, and the WD40 domain of DIC3 is docked onto Dyh3 HC. Within the HCs, there were also conformational changes. The Dyh3 showed more change in configuration with ∼90 degree rotation of the head compared with the inactive form whereas Dyh4 HC showed rather a compressing movement of the head side compared with Shulin-ODA structure (Fig. S5A and B). These conformational changes are mediated by hinge-like motions between the α-helical bundles (Fig. 3C). The center region (bundle 4-8) of Dyh3 showed a milder conformational change compared with other regions of the Dyh3 (Fig. 3C) since the LC tower is associated with this region. In detail, there is a slight change in conformation of both Dyh3 and LC tower (Fig. 3D and Fig. S5D). The conformational change was due to the association of α-helix bundle 8 of the Dyh3 with LC8/8d. This interaction was hindered in the Shulin-ODA complex since Shulin is keeping Dyh3’s bundle 8 away from LC8/8d. These tail rearrangements align the head domains to parallel configuration on the doublet (discussed later).

**Fig 3.**
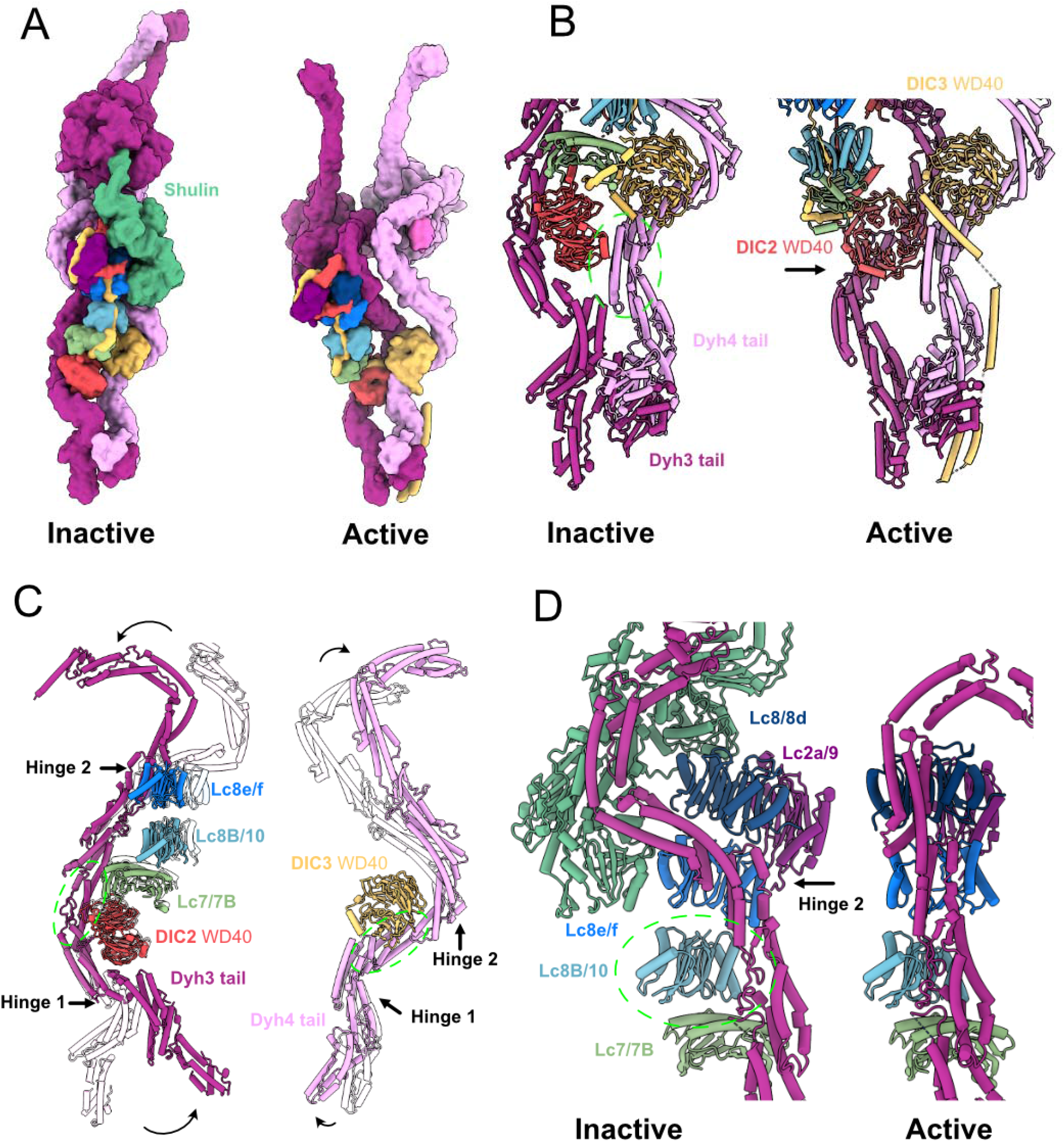
Comparison of the active ODA complex structure on the doublet and the inactive Shulin-ODA complex structure. (A) The surface rendering of the inactive ODA model (with Shulin bound) and the active ODA model. Dyh5 is not shown. (B) Remodeling of Dyh3 and Dyh4 tails and associated IC/LCs. The green circle indicates the region of alignment (Dyh4 residues 414-513, helix bundle 3). Dyh4 tail does not exhibit large conformational change while Dyh3 rotates almost 90 degrees, evident by the rotation of DIC2 WD40 domain. (C) Superimposition of Dyh3 and Dyh4 tails between the inactive (transparent) and the active ODA (Dyh3 is aligned based on res. 535-646, helix bundle 5 while Dyh4 is aligned on res. 414-513, helix bundle 4). The Dyh3 tail exhibits two hinges with larger rotations while the Dyh4 tail also exhibits two hinges with smaller rotations. (D) The changes in interaction between Dyh3 tail and the LC tower due to the remodeling of inactive to active conformations (two structures are aligned based on Lc8b/10). The conformations of Dyh3 bundle 8, Lc8/8d and Lc2a/9 are significantly different. The release of the Shulin leads to the rotation of the Dyh3 tail, which causes the different interaction pattern between Dyh3 and LC8/8d.

### Head domain structure of the ODA on the doublet

In the Shulin-ODA complex, the heads are fixed in the pre-powerstroke positions (Fig. 4A). In our active ODA complex structure, the linkers are in the post-powerstroke configuration both in the Dyh3 and Dyh4 heads. Therefore, the heads are ready to go through transition to next ATPase cycle with Shulin detached. Linker of the Dyh3 head was in a canonical post-powerstroke conformation (Fig. 4A). As for the Dyh4 head, LC3 was docked onto the AAA+ ring in the post-powerstroke configuration of the linker. LC3 was interacting with AAA4 of the binding position of the Dyh4 AAA+ ring. However, the binding scheme was different from that of Lis1 which also binds to the AAA3 of cytoplasmic dynein (Toropova, K., et al., 2014) (Fig. 4B). LC3 still interacts with AAA4 but it was closer to AAA3. LC1 was previously shown to bind to the MTBD of Dyh3 head (Ichikawa, M. et al., 2015, Toda, A., et al., 2020). LC3 is the second component found to be associated with the head domain of the ODA complex. Since LC3 was interacting with both linker and the AAA+ ring, we carefully examined the linker position of Dyh4. The linker was going at the middle of the apo state and the ADP state and slightly leaning toward AAA4 (Fig. 4C). This is a novel configuration of dynein linker. To perform powerstroke, LC3 is thought to be detached from the AAA+ ring (Fig. 4D).

**Fig 4.**
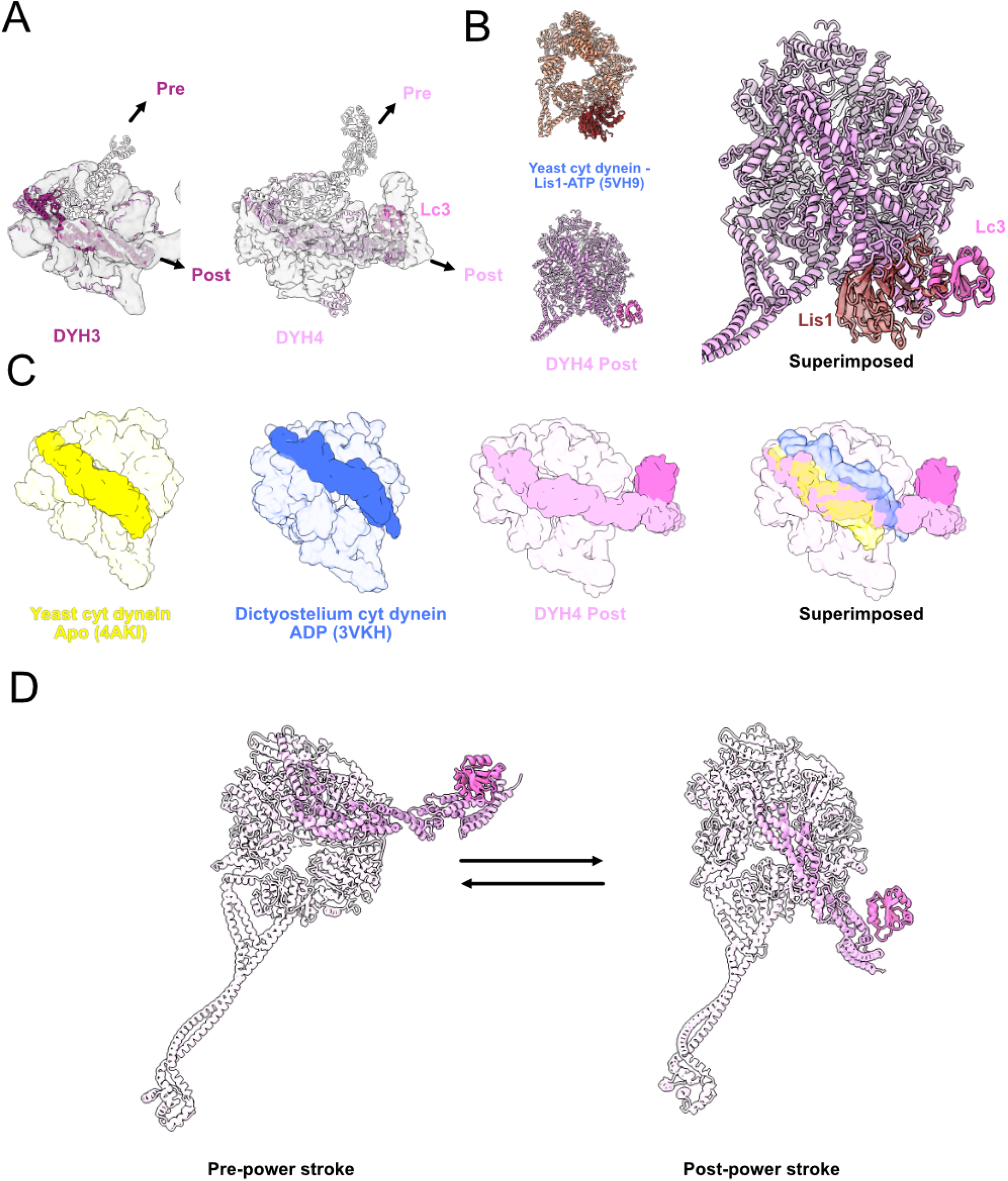
Structure of the head domain in ODA on the doublet. (A) Models of the Dyh3 and Dyh4 heads in post-powerstroke conformations are fitted in the cryo-EM maps of the ODA on the doublet. The Dyh3 and Dyh4 linkers in pre-powerstroke conformations in the inactive Shulin-ODA model are also superimposed (shown in transparent). Comparison of yeast cytoplasmic dynein bound to Lis1 (PDB: 5VH9) with the Dyh4 bound to LC3. LC3 also binds to the AAA+ ring but in an opposite site from Lis1. (C) Comparison of the linker of Dyh4 in post-powerstroke conformation with the yeast cytoplasmic dynein in apo state and Dictyostelium cytoplasmic dynein in ADP state reveals that the Dyh4 linker is positioned in between of the linker positions in apo and ADP states. The structures are aligned based on the AAA+ ring without linker. (D) Model for the conformation changes from pre-powerstroke to post-powerstrokes in Dyh4.

### Docking of the ODA complex to the DC induces the remodeling

How the regulatory protein Shulin detaches from the ODA complex? When Shulin-ODA structure was docked onto the doublet microtubule, the Shulin region did not show a steric clash (Fig. 5A). Therefore, we then examined the Shulin binding sites in the ODA complex on the doublet. Even though there was no other structure covering the Shulin binding sites, the regions involved in the interaction with Shulin were dispersed due to the rearrangement of the tail domain as well as the head domains (Fig. S5E). From these results, the remodeling of the ODA itself likely causes the detachment of Shulin. To understand the cause of the remodeling of the ODA complex, we focused on the region where ODA binds to the doublet. As mentioned above, the base region of the Dyh4 tail did not change significantly (Fig. 5A). In contrast, the tail region of Dyh3 remodeled and was located closer to the doublet (Fig. 5A and B). This remodeled region coincides with the part which interacts with DC3. In addition, the remodeled Dyh3 interacted with extended coiled-coil region of DC1/2 (Fig. 5C). The interaction between the head and the neighboring tail also thought to pull the Dyh3 HC.

**Fig 5.**
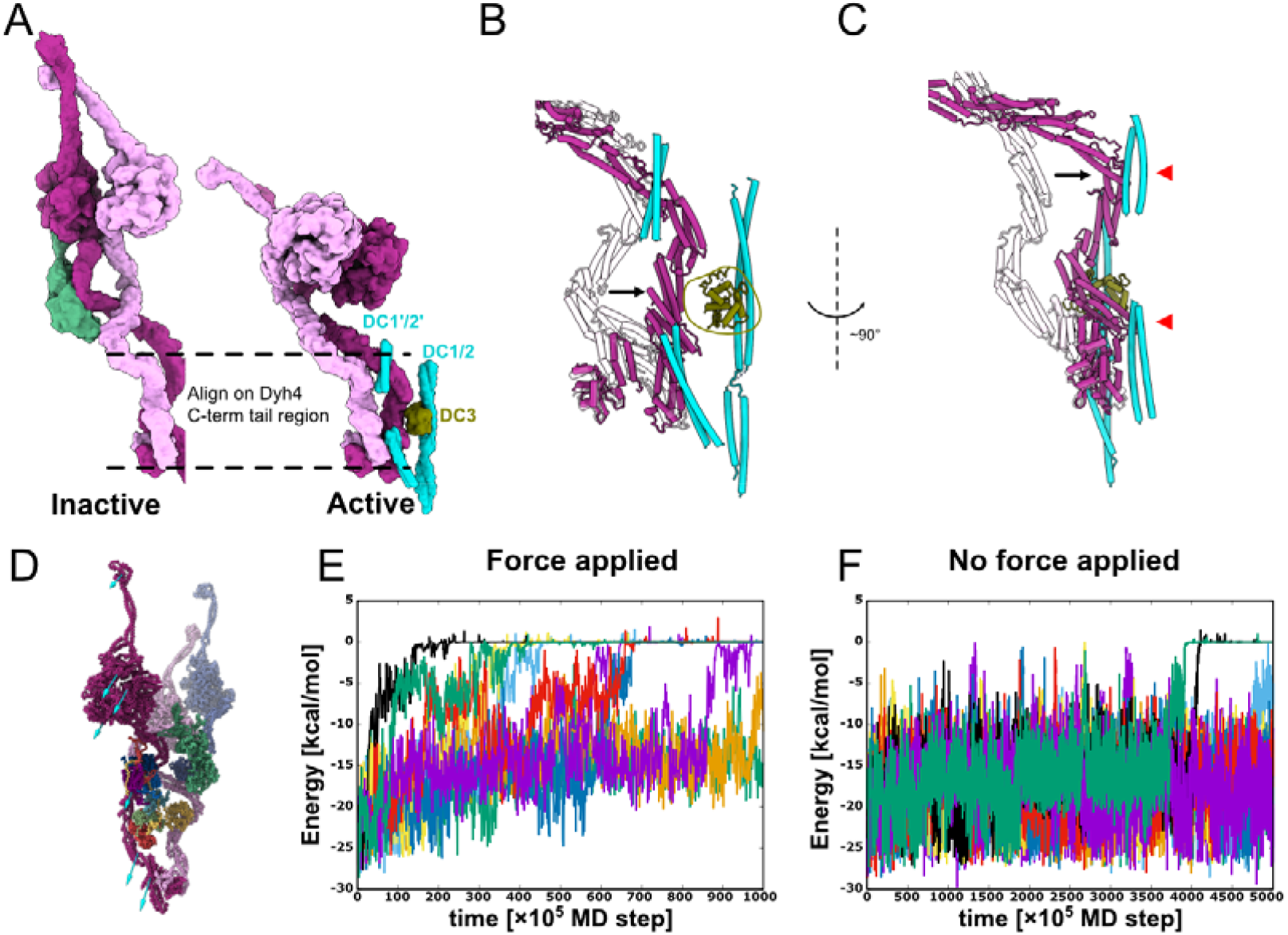
Remodeling of the tail domain and detachment of Shulin. (A) Models of the inactive and the active ODA. Dyh3 and Dyh4 parts from the models are shown with Shulin and docking complex, respectively. The two structures are aligned based on the Dyh4 residues 414-513 (helix bundle 3). (B, C) Detailed comparison of the Dyh3 before and after the remodeling. With the two structures aligned as in A, Dyh3 tail in the active conformation indicates significant shift towards DC3 (B). Similarly, the Dyh3 tail in the active conformation indicates a significant shift toward the extended coiled-coil region of DC1/2 (∼320-400 aa). Red arrowheads indicate the interaction sites of Dyh3 with the extended coiled-coil region of DC1/2. (D). Coarse grain molecular dynamics of the inactive Shulin-ODA complex. External force was applied in the direction of the arrow to simulate the remodeling of Dyh3 upon binding to the doublet microtubule. (E, F) Energy trajectories between Shulin and the ODA complex with (E) and without (F) external force applied to Dyh3. 8 out of 10 setups showed detachment of Shulin (energy = 0) with external force in 1,000 × 10^5^ MD steps (E). In contrast, only 2 out of 10 trajectories showed Shulin detachment without applied force even with 5,000 × 10^5^ MD steps (F).

To test if the remodeling of the Dyh3 HC causes the detachment of Shulin, we performed a MD simulation. To mimic the remodeling of Dyh3, external force was applied to Dyh3 of the Shulin-ODA complex (Fig. 5D, see Materials and methods for the details). In eight out of ten MD trajectories, Shulin detached within 1,000 frames (Fig. 5E). Without external force applied, only two out of ten trajectories showed detachment within 5,000 frames, which is a five times longer time frame than the case with external force (Fig. 5F). Therefore, the attraction by DCs and the adjacent ODA is likely to be important for Shulin detachment.

## Discussion

Our structure presents a physiological binding state of the ODA to the doublet via the DC complex in an unprecedented resolution. The ODA complex is docked onto the doublet by two-sites via DCs. In addition, the ODA interacts with the neighboring ODA complex. These interactions are crucial for the stable attachment of the ODA to the doublet, and therefore, force generation by the ODA complex within cilia. The ODA is bottomed up by DC “heals” so that it can associate with the neighboring doublet properly. In different species, DCs might tune the ODA interaction schemes. DC3 in Tetrahymena and Chlamydomonas contains several EF-hand motifs. We speculate that the interaction of DC3 to the ODA can change based on the intracellular calcium ion concentration, leading to the regulation of ODA activity. In Chlamydomonas, high concentration of calcium ions is proposed to induce a symmetric waveform instead of the normal asymmetric breaststroke waveform (Bessen, M. et al., 1980). DC3 is not conserved in vertebrates. In humans, the DC assembly affects CCDC114, CCDC151 and ARMC4 (Hjeij, R. et al., 2014). Therefore, ARMC4 might act as DC3 in humans. However, ARMC4 (1,044 aa) is a significantly larger protein than DC3 (309 aa in Tetrahymena). Comparing the tomography maps of human cilia and Tetrahymena, the human ODA also docks on the doublet at two sites (Fig. S4B) (Stoddard et al., 2018; Lin et al., 2014a). However, both docking sites are different from the Tetrahymena ODA. The docking site close to Tetrahymena DC3 docking site is shifted (Fig. S4B), while the other docking site attaches to PF-A7, instead of attaching to the DC1/2 between PFs-A7 and A8 as is the case in Tetrahymena. In humans, mutations leading to early termination or frameshift of CCDC151 (p.Glu109X, p.Glu309X and p.Ser419X) and CCDC114 (p.Ala248Serfs52 and p.Ser432Argfs*7) cause primary ciliary dyskinesia (Hjeij, R. et al., 2014; Knowles, M. R. et al., 2013; Zhang, W. et al., 2019) (Fig. S4A). These mutations map to both the regions of the DC1/2 of Tetrahymena binding site to the doublet and also binding to the Dyh3 tails as proposed by our structures. This implies the importance of the extended coiled-coil region of DC1/2 for the assembly of the ODA and for the remodeling of the tDA.

By comparing the active ODA complex structure with the recently obtained inactive form of the ODA complex, we were able to reveal the details of the remodeling of the ODA complex. The observed docking of the LC3 to the AAA+ ring of Dyh4 in the active ODA complex was surprising. Previous cryo-ET studies revealed that the βHC of the sea urchin sperm (equivalent to the Dyh4 of Tetrahymena) can undergo an ATPase cycle (Lin, J. et al., 2014b). Therefore, in the ATP cycle of the Dyh4, the linker detaches from and re-attaches to the AAA+ ring (Fig. 4D). LC3 is a thioredoxin component which was proposed to regulate the ODA activity in response to redox state of the cilia (Wakabayashi, K. and King, S, M., 2006). Since LC3 was shown to interact with other proteins within the cilia depending on the redox state, LC3 might recruit other regulatory proteins to the Dyh4 AAA+ ring to modulate its motor activity.

Based on our structure and MD simulation, we present a model of how the ODA attachment induces remodeling and activation of the ODA complex (Fig. 6). First, the Shulin-ODA complex docks onto the doublet using DC3 as a marker (Fig. 6A). Next, the Dyh3 tail region starts to remodel. The extended coiled-coil region of DC1/2 catches the Dyh3 tail domain and facilitates the remodeling of the Dyh3 tail region (Fig. 6B). Due to the remodeling in the Dyh3 tail region, the binding of Shulin becomes destabilized. After Shulin detaches, the remodeling of the ODA complex is propagated toward the heads (Fig. 6C). Without Shulin, the head domains would be released to be in a parallel configuration. Furthermore, the linker would be in post-powerstroke configurations and ready to exert force in cilia. Interaction of the head domain and the neighboring ODA complex would enhance this process. Our model illustrates a simple yet robust activation mechanisms of the ODA complex which does not require other external factors.

**Fig 6.**
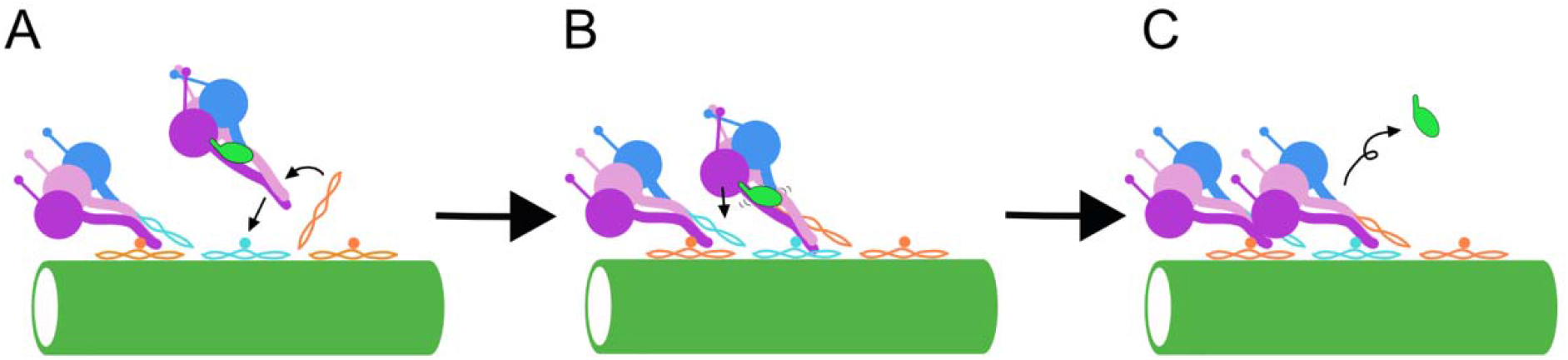
Model of the activation mechanism of the ODA. (A) Shulin-ODA docks on the doublet through interaction between the tail region of Dyh3 and the DC3. (B) Extended coiled-coil region of the DC1/2 catches the ODA complex at the tail domain and facilitates the remodeling of the tail. Head domain of Dyh3 also interacts with the neighboring ODA complex’s tail. Therefore, the Dyh3 is “pulled” toward the doublet and binding of the Shulin destabilizes. (C) Shulin detaches from the ODA complex and the head domains are remodeled and aligned parallel to each other. Head domains are also in post-powerstroke conformations and ready to exert force.

The remodeling of the tail involves an interaction with the extended coiled-coil part of DC1/2. This is the reminiscence of the cytoplasmic dynein’s case. Cytoplasmic dynein-1 takes inactive conformation with two heads stacked together (Torisawa T., et al., 2014), when it binds to dynactin and adaptor proteins, such as BICD2, BICDR1 and HOOK3, the tail domain remodels and the two heads becomes separated (Zhang, K., et al., 2017; Urnavicius, L. et al., 2018). These adaptor proteins are also coiled-coil proteins and interact with the cytoplasmic dynein heavy chain by two sites to stabilize the remodeled conformation. In the cytoplasmic dynein, one of the ICs’ WD domain interacts with the other remodeled HC similarly observed in the ODA’s ICs. Thus, similar mechanisms are exploited for cytoplasmic dynein-1 and the ODA for the activation. The stacking of the head is also conserved with cytoplasmic dynein-2 (IFT dynein) (Toropova, K. et al., 2017; Toropova, K. et al., 2019). Thus, it would be interesting to clarify if there are similar adaptor proteins to remodel the tail when dynein-2 is working in the retrograde intraflagellar transport. The heterodimeric inner arm dynein-f has different subunit components and head arrangement compared with the ODA (Heuser, T. et al., 2012). Therefore, future study is needed to show whether the same mechanism is exploited with two-headed axonemal dynein-f when incorporated to the doublet.

In conclusion, we have shown the ODA complex structure on the doublet by cryo-EM and obtained insights into how the ODA is activated inside the cilia. Our findings would be the important foundation to understand the generation of ciliary structure.

## Materials & Methods

### Growth of *Tetrahymena* cells for isolation

1 ml of the saturated Tetrahymena cells in SPP media (Gorovsky *et al*., 1975) was transferred into 50 ml liquid SPP and grown overnight with shaking at 150 rpm at 30□ in the shaker incubator (ThermoFisher MAXQ8000). The overnight 50 ml culture was added to 900 ml of SPP and grown for approximately two days with shaking at 150 rpm at 30□ (MAXQ8000) until the cell density reached an OD at 600 nm of 0.7.

### Cilia isolation by dibucaine treatment

The *Tetrahymena* culture was centrifuged at 700g for 10 min at 22°C with slow deceleration (Avanti, Rotor JLA-8.1000). The cell pellet was resuspended with 10 ml of SPP and the volume was adjusted to 24 ml. The cell suspension was transferred to a 250 ml Erlenmeyer flask on ice. 1 ml dibucaine in SPP at 25 mg/ml was added to the flask to make a final concentration of 1 mg/ml, and the cell suspension was gently swirled over ice for exactly 60 seconds. 75 ml of ice-cold liquid SPP media was immediately added, and the 100 ml cilia suspension was then split into two 50 ml conical tubes. The cell suspension was centrifuged at 2,000g for 10 min at 4°C with slow deceleration (Sorvall ST 16R, Rotor 75003181). The cilia-containing supernatant was transferred to 4 centrifuge tubes for the Beckman Coulter JA 25.50 rotor, approximately 20 ml per tube. The tubes containing the cilia-containing supernatant were centrifuged at 17,000g for 40 min at 4°C with slow deceleration (Avanti, Rotor JA25.50). The supernatants were discarded and the pellets were carefully washed with theice-cold Cilia Wash Buffer (50 mM HEPES at pH 7.4, 3 mM MgSO4, 0.1 mM EGTA, 1 mM DTT, 250 mM sucrose), before resuspension in 250 uL of the ice cold Cilia Wash Buffer. The cilia were pelleted by centrifugation with 16,000xg for 10 min at 4°C in a microfuge, flash-frozen with liquid nitrogen and stored in the -80°C.

### Purification of intact doublet microtubule

The thawed cilia pellet was resuspended in 250 μl ice-cold Cilia Final Buffer (50 mM HEPES at pH 7.4, 3 mM MgSO4, 0.1 mM EGTA, 1 mM DTT, 0.5% Trehalose, 1 mM PMSF). The concentration of total protein was measured using the Bradford Assay. The resuspended cilia sample was centrifuged at 16,000g for 10 min at 4°C in a microfuge (Eppendorf, Centrifuge 5415D). The supernatant was discarded, and the pellet was resuspended once more with 250 μl Cilia Final Buffer. NP-40 alternative (Millipore Sigma, 492016) was added to the final concentration of 15% and the total solution was resuspended. The samples were incubated on ice for 30 min before the demembraned cilia were centrifuged at 16,000g for 10 min at 4°C in a microfuge (Eppendorf, Centrifuge 5415D). The supernatant was discarded, and the axoneme-containing pellet was resuspended to 245 μl of Cilia Final Buffer. Final concentration of 1 mM ADP was added to the axonemes and incubated at room temperature for 10 minutes. Then, final concentration of 0.1 mM ATP was added to the solution and incubated at the room temperature for 10 minutes.

### Cryo-EM sample preparation

Wild-type, K40R and MEC17-KO doublet concentrations were adjusted to 3 mg/ml. Quantifoil R2/2 (Electron Microscopy Sciences, #Q2100CR2) were treated with chloroform overnight and then glow discharged (30 seconds at 25 mAh). 3.5 μl of axoneme were applied to the pretreated grids and bolotted by the Vitrobot Mark IV using blot force 3, blot time of 5 seconds, and drain time of 0.5 seconds.

### Cryo-EM data acquisition

Movies of the doublet microtubules were acquired at 64kx nominal magnification (calculated pixel size of 1.370 Å/pixel) with counting mode using Titan Krios 300 kV FEG electron microscope (Thermo Fisher Scientific) with the direct electron detector K3 Summit (Gatan, Inc.) and the Bioquantum energy filter (Gatan, Inc.). The data were acquired using SerialEM (Mastronarde, D. N., 2005). The WT and MEC-17 datasets were collected using a total dose of 45 electrons per Å^2^ over 40 frames. The K40R dataset was with a total dose of 73 electrons per Å^2^ per frame over 30 frames. The defocus range was between -1.0 and -3.0 μm.

### Cryo-EM image processing

The movies were motion corrected and dose-weighted using MotionCor2 (Zheng, S. Q. et al., 2017) implemented in Relion3 (Zivanov, J. et al., 2018) and the contrast transfer function parameters were estimated using Gctf (Zhang, K., 2016). After discarding micrographs with apparent drift and ice contamination, bad contrast transfer function estimation (3,074, 2,410, 4,283 micrographs for WT, K40R and MEC17 data, respectively). K40R and MEC17 data were merged since the ODA structures were the same in both mutants (Yang, S. et al., manuscript in preparation). The filaments were picked using e2helixboxer (Tang, G. et al., 2007).

The particles of 512 x 512 pixels were initially picked with 8-nm periodicity, binned twice and pre-aligned using a modified version of the Iterative Helical Real Space Reconstruction script (Egelman, E. H., 2007) in SPIDER (Frank, J. et al., 1996) to work with non-helical symmetry. After that, the alignment parameters were transferred to Frealign and the particles were aligned by Frealign for 6 iterations (Grigorieff, N., 2007). The alignment parameters were then converted into Relion 3.0. The particles were scaled to 1.744□Å/pixel (box size 402 pixel) for image processing to reduce memory usage. The 8-nm particles then underwent iterative per-particles-defocus refinement and Bayesian polishing in Relion 3.

The signal of the tubulin lattice was subtracted from each particle. The subtracted particles were subjected to 3D classification into three classes to obtain the 24-nm repeat particles with the ODA in the center. The 24-nm particles were then refined, resulting in resolution of 4.3 and 3.9 Å for WT, and K40R & MEC17 combined data from 71,462 and 139,548 particles, respectively. At this point, it is clear that the WT doublet did not retain a lot of ODA, only the docking complex (Fig. S1D and E). Therefore, we continued the next step with the K40R and MEC17 combined dataset.

The 24-nm particles were boxed out at a box size of 804 pixel at 1.744□Å/pixel and underwent Bayesian polishing. The ODA are then centered and boxed out of the imaging using the Localized Reconstruction in Scipion (De la Rosa-Trevín, J. M. et al., 2016; Abrishami, V. et al., 2020). The particles containing the ODA were then refined using Relion3 and cryoSPARC (Zivanov, J. et al., 2018; Punjani, A. et al., 2017). We first refined a single entire ODA region (Fig. S2A). Then, we refined the ODA tail and head regions separately using focus refinement with masks for different regions (Fig. S2B). The resolutions for different regions of the ODA are in the range of 5.5 to 7Å (Fig. S2B-D).

### Model building of the ODA complex on the doublet

First, the Shulin-ODA complex model was separated into two regions: (1) Dyh3 & IC/LC tower region and (2) Dyh4, 5 & DIC3’s WD domain. These two regions were rigid-body fitted into the active ODA map based on the tail region. Then, ICs and LCs were rigid-body fitted one by one to our cryo-EM map. Dyh3 and Dyh4 were segmented into several regions and further fitted to the density. Since the IC and LC regions were fitted quite well to our map at this point, the density corresponding to ICs and LCs were subtracted from the whole tail map and then the HC regions were fitted to the map more precisely. For the head part, the whole head domains were fitted to our cryo-EM map. We noticed that the linker parts of the Shulin-ODA complex were sticking out since our maps are in post-powerstroke conformations. Thus, the linker part was segmented and fitted separately. For Dyh4, the linker region and LC3 were fitted together to our map and then fitted further separately. Finally, the model was integrated, and refined by Coot (Emsley, P. et al., 2010) in our map and real space refined using Phenix (Adams, P. D. et al., 2010).

### Docking Complex Modelling

DC2 (Q233H6) and DC3 (I7M2C6) of Tetrahymena were found by BLASTP from the Chlamydomonas DC2 (A8JF70) and DC3 (Q7Y0H2). Their initial models were constructed by MODELLER (Ŝali, A. and Blundell., T. L., 1993) from the DC2 and DC3 structure (PDB 6U42). Q233H6 and I7M2C6 were modelled by Coot (Emsley, P. et al., 2010) in the density of the 24-nm average doublet microtubule structure (Fig. S3A), in which the docking complex density is much clearer.

DC1 from Chlamydomonas (A8IPZ5) did not yield a homolog in Tetrahymena using BLAST. Since we could clearly see a density at the place of DC1 in the density map and DC1 from *Chlamydomonas* is not conserved in humans, we reasoned that there is another coiled coil protein in place of DC1. In humans, CCDC151 is known to be important for the docking complex and causes ciliopathy similar to CCDC114 (human DC2 homolog) (Hjeij, R. et al., 2014; Zhang, W. et al., 2019). The human CCDC151 localizes to respiratory ciliary axonemes and interacts with CCDC114 (DC2 homolog). The proposed homolog of CCDC151 in Chlamydomonas is ODA10. While this protein is important for the ODA assembly in Chlamydomonas (Dean, A. B. and Mitchell, D. R., 2013), it does not seem to localize in the axoneme like DC1/2 from the MS pattern (Pazour, G. J. et al., 2005).

When we blasted CCDC151, we found Q22T00 as a homolog. To see the MS profile of Q22T00, MS analyses were performed for untreated and salt-treated doublet samples as described in Dai, D. et al., (2020). The MS profiles of Q22T00 were similar to DC2 and DC3 (Table S1). The coiled coil prediction of Q22T00 using COILS program (Lupas, A. et al., 1991) also coincided with Chlamydomonas DC1 (Fig. S3B). Therefore, we concluded that despite the weak conservation, Q22T00 is the DC1 in *Tetrahymena*. We constructed the initial model of Q22T00 to the DC1 using MODELLER. Q22T00 was then modelled by Coot in the density of the 24-nm average doublet microtubule structure (Fig. S3A), in which the docking complex density is much clearer. The secondary prediction of Q22T00 fitted well with helical density observed in the density map. For the extended coiled-coil part of DC1/2, poly-alanine model was generated based on two regions of the coiled-coil of BICDR1 model (105-161 and 202-245 aa from PDB: 6F1T) and fitted to our cryo-EM map.

### Visualization

The maps and models are visualized using ChimeraX (Goddard, T. D. et al., 2018).

### MD simulation

The inactive Shulin-ODA structure had some missing residues. For MD simulation, we modelled loops for missing regions by MODELLER (Ŝali, A. and Blundell., T. L., 1993). At the same time, two intermediated parts of Dyh5 (531-636, 3,270-3,382), and both ends of the Dic2 and Dic3 sequences (1-74, 636-667, and 1-17, 511-670, respectively) were removed. Then, we did a coarse-grained MD simulation using this model. In the model, 1 amino-acid was treated as 1 bead and each bead located at the Cα position. The force field for observing dynamics used AICG2+ (Li, W., et al., 2012; Li, W. et al., 2014). In the AICG2+, the reference structure was assumed as the most stable conformation, and their parameters are defined from the reference of all-atom structures. Recently, AICG2+ represented the conformational change of cytoplasmic dynein, and it clarified the allosteric conformational change cascade of dynein’s motor domain (Kubo, S. et al., 2017). Here, by comparing the inactive Shulin-ODA model with the active ODA structure on the doublet, we identified chains that keep their binding and chains which change their binding schemes. Based on this information, we adjusted the parameters that determine the attraction between the chains. First, since Dyh3, 4, and 5 clearly have a different contact style, we set the attraction force between these chains to be 0.5 times of the default. Next, since the contacts between Dyh3, Dic2, and the LC tower were not significantly changed, we increased the attraction by a factor of 10 in order to treat this chain as a rigid body. Also, since there was no significant difference between Dyh4 and Dic3 contacts, we increased the force of attraction by a factor of 10. Lastly, the attractive force between Shulin and others was decreased by 0.3 times the default in order to observe the Shulin detachment in a reasonable simulation time. After that, we run the simulations on several systems. We performed 10 simulations for all systems using CafeMol version 2.1 (Kenzaki, H. et al., 2011). If there is not written additional, each MD simulation took 5 × 10^7^ MD steps, and they were conducted by the underdamped Langevin dynamics at 300K temperature. We set the friction coefficient to 0.02 (CafeMol unit) and default values were used for others.

## Acknowledgements

We thank Drs. Kaustuv Basu, Kelly Sears, Mike Strauss, Ferdos Abid Ali and Andrew P. Carter for helping with data collection, processing and modelling. This work was supported by JST, PRESTO Grant Number JPMJPR20E1, JSPS KAKENHI Grant Numbers JP19K23726 and JP20K15733, and the Foundation for Nara Institute of Science and Technology (R2290001) to MI. SK is supported by JSPS Overseas Research Fellowships. KHB is supported by the grants from Canadian Institutes of Health Research (PJT-156354), Natural Sciences and Engineering Research Council of Canada (RGPIN-2016-04954) and Canadian Institute for Advanced Research (Arzieli Global Scholar Program).

## Conflicts of Interest

The authors declare no conflicts of interest.

## Author Contributions

KHB and MI conceived the project and designed the experiments. JG generated mutant Tetrahymena strains. SKY and CB performed culture of the cells, purification of microtubule fractions from cilia with the help of DD and MV. SKY and CB performed vitrification of the grids, collected data. SKY, SK and KHB performed the cryo-EM data analysis of the doublet. MI generated a model of the ODA complex on the doublet. SK performed the MD simulation. SK, MI and KHB interpreted the structure. All authors were involved in the manuscript writing process.

## Supplementary Figures

**Supplementary Figure 1.**
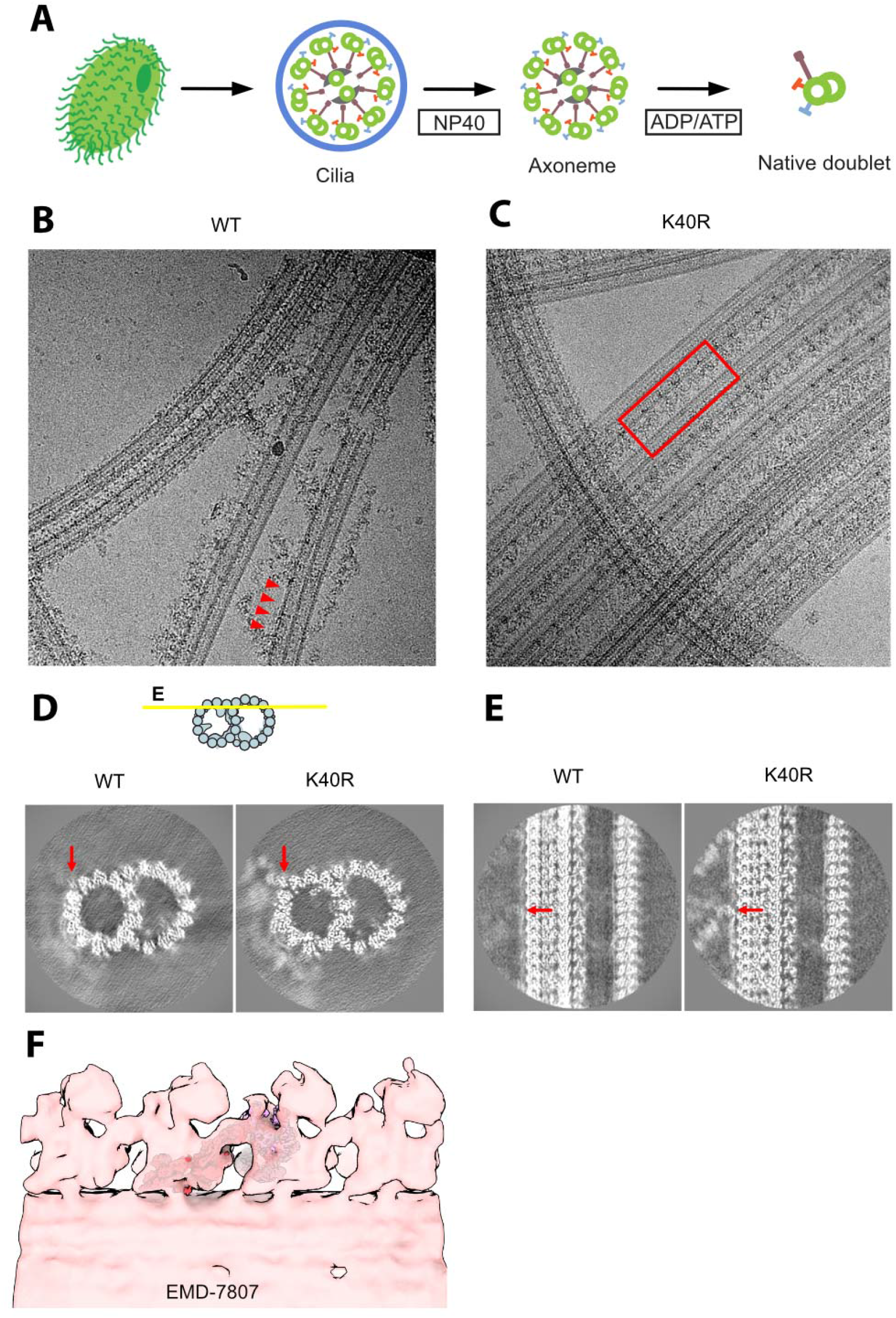
Sample preparation and cryo-EM of the doublet microtubules. (A) The schematic of our isolation strategy for the intact doublet microtubules. (B, C) Micrographs of doublet microtubules from *Tetrahymena* WT (B) and K40R mutant (C). The red arrowheads indicate the ODA complex falling off from the doublet. The red rectangle indicates the row of intact ODA in the K40R mutant. (D, E) 24 nm-structure of doublet from WT and K40R showing the DC is intact in both cases while ODA is clearly present only in K40R. Red arrows indicate the docking complex. (F) Fitting of our high-resolution structure into the tomographic map of Tetrahymena showing it is physiological (ΔRib72B mutant rescued with Rib72B-GFP) (EMD-7807, Stoddard, D. et al., 2018).

**Supplementary Figure 2.**
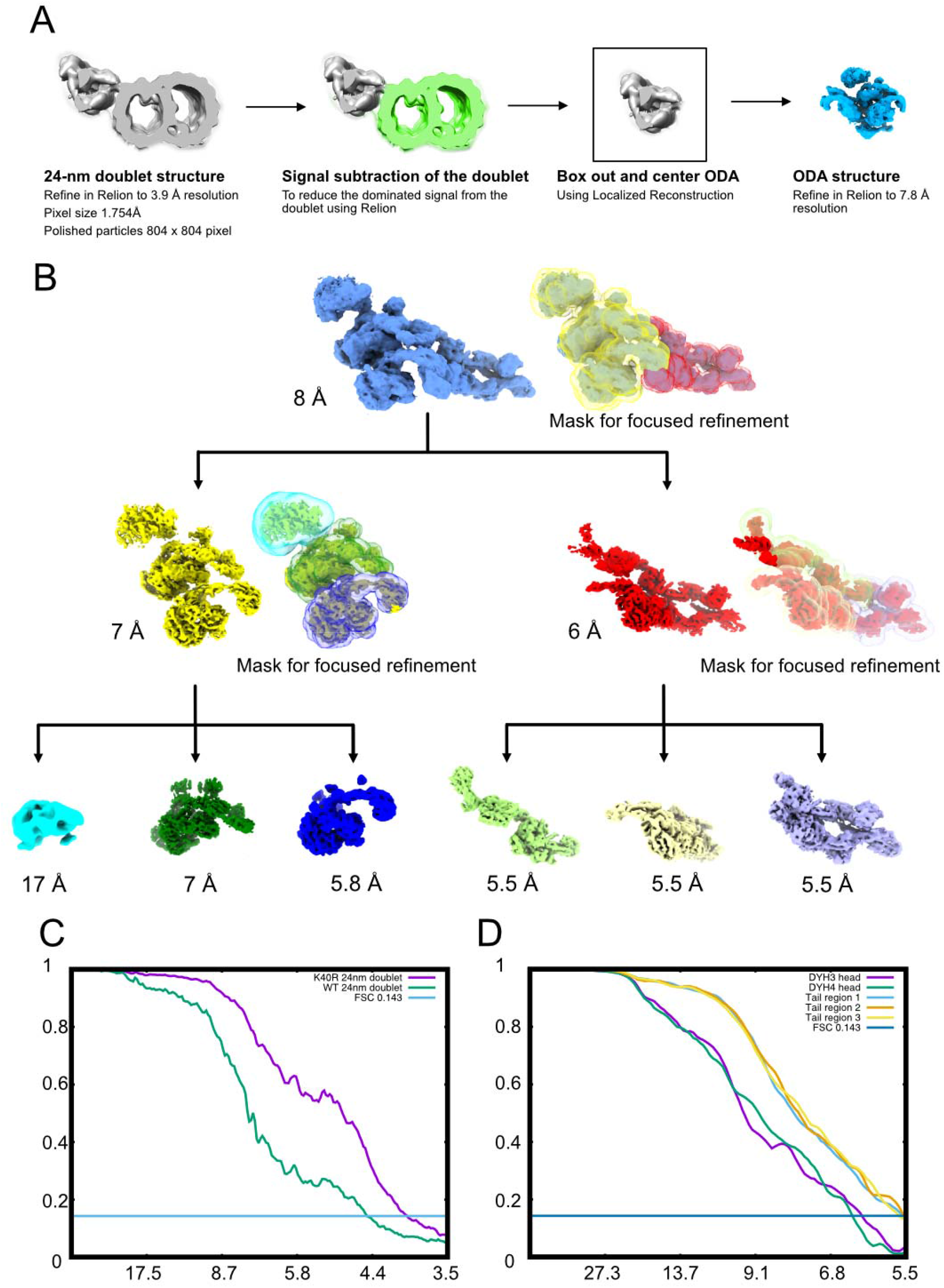
Cryo-EM processing strategy. (A) Alignment strategy for ODA particles. First, we obtained the 24-nm doublet structures. After that, we performed signal subtraction of the doublet microtubule. We centered and boxed out the ODA particles and performed refinement of the entire ODA particles. (B) Focus refinement strategy for different regions of the ODA complex. The Dyh5 seems to be too flexible, therefore we can only obtain Dyh4 head at 17 Å resolution. (C) Fourier Shell Correlation of the doublet of WT and K40R & MEC17 combined data. (D) Fourier Shell Correlation of the different regions of the ODA complex by focus refinement.

**Supplementary Figure 3.**
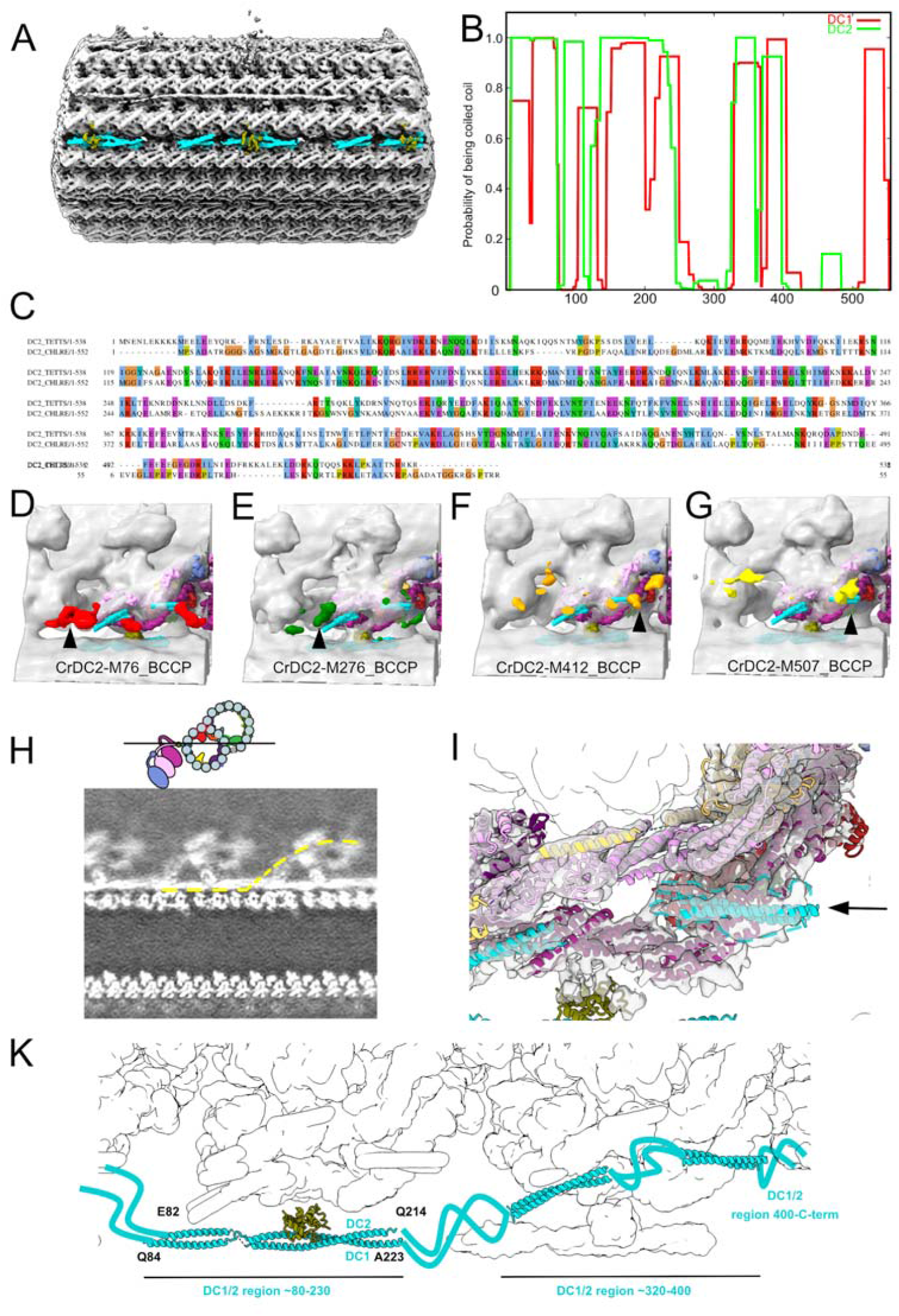
Modelling of the Docking Complex. (A) The DC density on the doublet microtubule between PFs-A7 and A8. (B) Prediction of coiled coil for DC1 and DC2 using COILS with a window size of 28 (Lupas, A. et al., 1991). (C) Sequence alignment of Chlamydomonas DC1 and Tetrahymena CCDC151 homolog (Q22T00). (D, E, F, G) Structures of Chlamydomonas ODA reconstituted on microtubules with Biotin carboxyl carrier protein (BCCP) tagged in different regions of DC2 (residue 76, 276, 412 and 507) from Oda, T. et al., (2016) (EMD-6508, 6509, 6510, 6511). The enhanced signals of BCCP-tag are indicated in colors. (H) Slice from a density map showing the docking complex (position indicated in the cartoon). The yellow line indicates one continuous DC1/2. (I). The globular density at the end of the coiled coil of DC1/2 (black arrow and dotted line). (K). The model of the DC based on our analysis.

**Supplementary Figure 4.**
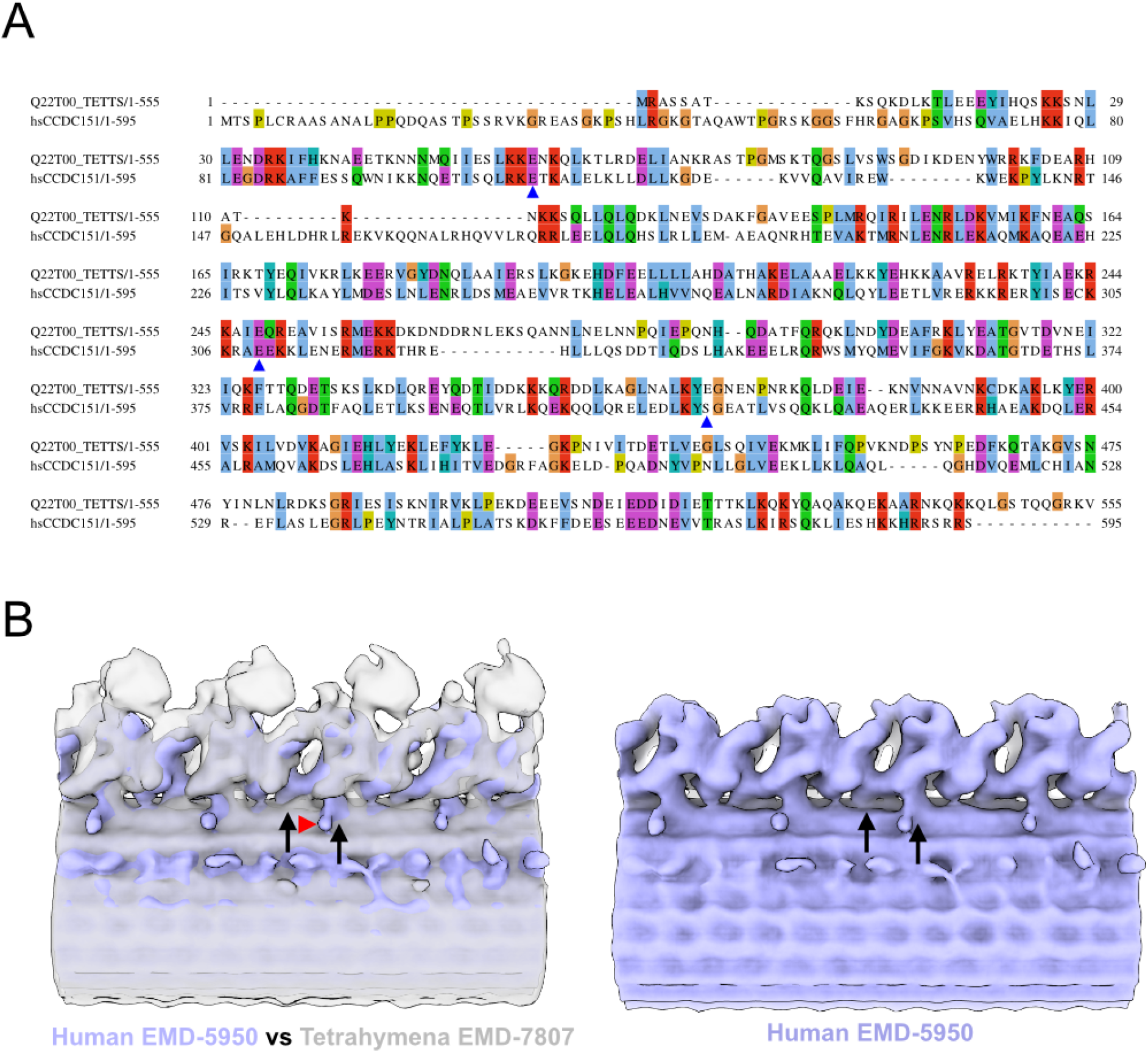
Data related to Docking complex. (A) Sequence alignment of Tetrahymena DC1 (Q22T00_TETTS) and human CCDC151. Blue arrowhead indicates the residues at which early terminations are found in patients with ciliopathy (Hjeij, R. et al., 2014). (B) Overlapping of the tomographic structure of the axonemes from Tetrahymena (EMD-7807, Stoddard, D. et al., 2018) and human (EMD-5950, Lin, J. et al., 2014) (left panel) and the human axoneme alone (right). The two maps are aligned based on the radial spokes on the opposite sides. It is clear that the docking point of the human ODA (black arrows) are different from Tetrahymena ODA. In addition, there is an extra density protruding from the docking complex of humans (red arrowhead).

**Supplementary Figure 5:**
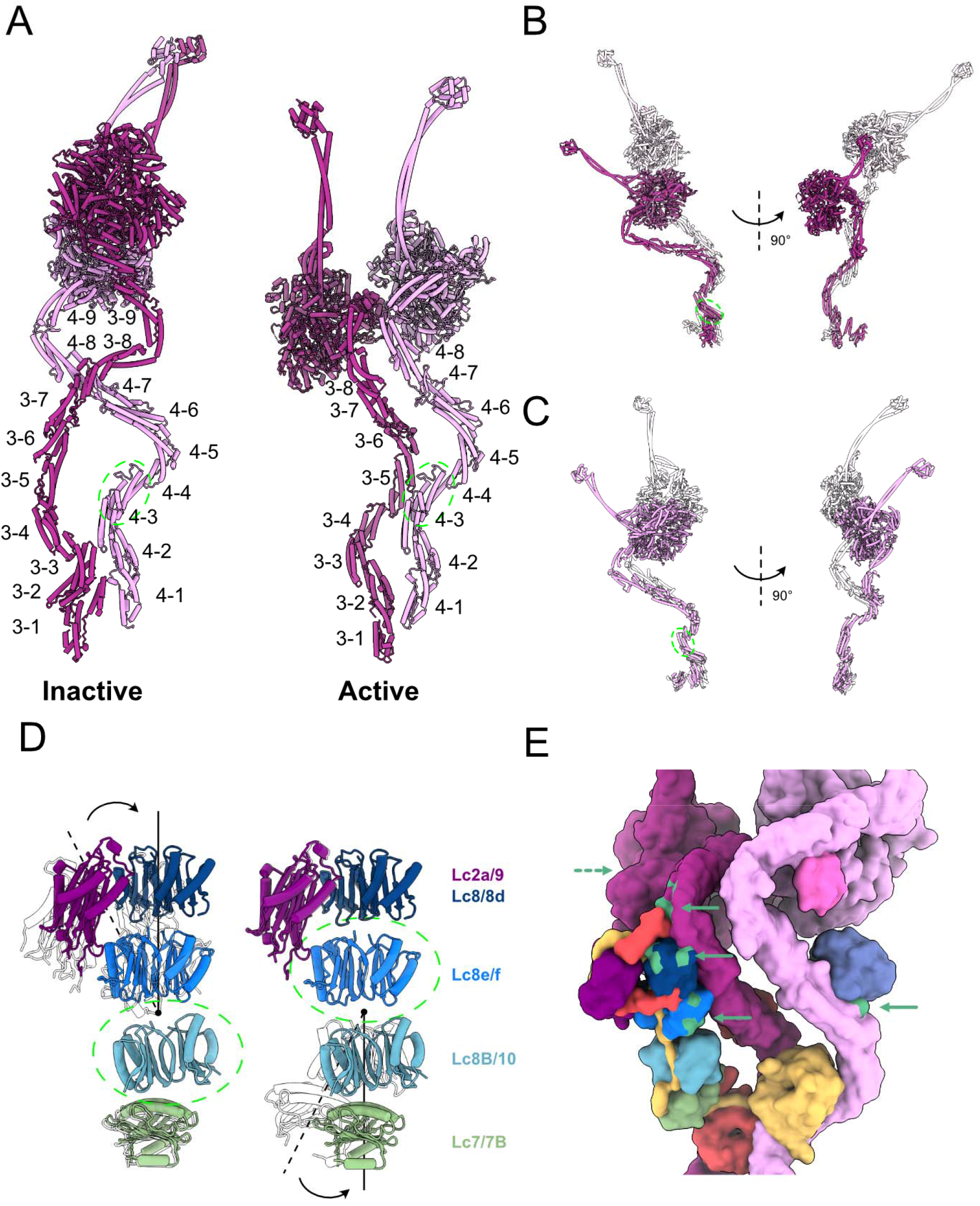
Data related to the remodeling of the ODA complex. (A) The labeling of helix bundles in the inactive and active Dyh3 and Dyh4. Inactive and active structures are aligned on helix bundle 4 of Dyh4 (residues 414-513). (B) Alignment of the inactive and active Dyh3 at helix bundle 3 (residue 448-536) showing ∼90 degree rotation of the head domain (top). Alignment of inactive and active Dyh4 at helix bundle 3 (residue 414-513) showing compressing conformational changes (bottom). (C) Bending conformational change of the LC tower. LC tower from the active ODA complex is in colors and the LC-tower from the Shulin-ODA is shown in transparent. LC towers are aligned based on either Lc8B/10 (left) or Lc8e/f (right) as indicated by green dashed circles. (D) Bending conformational change of the LC tower. LC tower from the active ODA complex is in colors and the LC-tower from the Shulin-ODA is shown in transparent. LC towers are aligned based on either Lc8B/10 (left) or Lc8e/f (right) as indicated by green dashed circles. (E) Regions of the ODA that interact with Shulin in the inactive conformation (green regions with green arrowhead) are spread out in the active conformation. The dash arrow indicates the region of Dyh3 head interacting with C3 domain of Shulin, now at the back of the view.

## Supplementary Table

**Supplementary Table 1:** MS total spectrum count of DC1, DC2 and DC3 from Tetrahymena WT doublet using native purification and NaCl treatment. In the NaCl treated doublet, most of the DC components were washed out.

| Name | Name | UniprotID | K40R | WT | WT (NaCl) |
| --- | --- | --- | --- | --- | --- |
| Heavy chain | Dyh3 | Q22A67 | 239,232,229 | 263,250,245 | 1,0,0 |
|  | Dyh4 | I7M9J2 | 259,241,247 | 264,255,240 | 0,0,0 |
|  | Dyh5 | I7M6H4 | 223,215,199 | 239,241,218 | 2,2,0 |
| Intermediate chain | Dic2 | I7M008 | 36,32,31 | 38,37,32 | 6,10,9 |
|  | Dic3 | Q23FU1 | 40,38,37 | 38,39,35 | 15,11,12 |
| Light chain | Lc3 | A4VD75 | 9,9,7 | 7,7,6 | 0,0,0 |
|  | Lc4a | Q22C78 | 5,5,3 | 3,3,2 | 0,0,0 |
|  | Lc1 | I7M1N7 | 22,22,20 | 19,17,18 | 0,0,0 |
|  | Lc2a | Q1HGH8 | 4,4,4 | 5,5,5 | 1,3,0 |
|  | Lc9 | A4VEB3 | 4,3,3 | 5,5,3 | 0,0,0 |
|  | Lc8 | W7XJB1 | 5,4,4 | 3,4,3 | 0,0,0 |
|  | Lc8d | Q24CE5 | 7,6,5 | 7,7,5 | 0,0,0 |
|  | Lc8e | Q24DI9 | 3,2,3 | 1,3,2 | 0,0,0 |
|  | Lc8f | Q22R86 | 4,4,5 | 3,4,5 | 0,0,0 |
|  | Lc10 | I7MCM8 | 2,0,2 | 0,2,0 | 0,0,0 |
|  | Lc8b | A4VE64 | 6,6,6 | 2,2,0 | 0,0,0 |
|  | Lc7 | I7MHB1 | 8,8,8 | 6,7,7 | 0,3,3 |
|  | Lc7b | Q1HFX1 | 6,4,5 | 5,5,4 | 0,0,0 |
|  | Lc7a* | Q1HFX2 | 4,4,4 | 3,3,3 | 0,0,0 |
|  | Lc2b* | I7M0N9 | 4,6,6 | 4,4,4 | 0,0,0 |
|  | Tct1a* | I7MGG2 | 0,0,0 | 0,0,0 | 0,0,0 |
|  | Tct1b* | Q231T5 | 2,2,2 | 0,0,0 | 0,0,0 |
| Docking Complex | DC1 | Q22T00 | 29,26,28 | 27,26,22 | 0,0,2 |
|  | DC2 | Q233H6 | 20,19,17 | 24,25,24 | 0,3,2 |
|  | DC3 | I7M2C6 | 15,13,11 | 17,16,17 | 4,3,4 |

\* Components do not present in ODA according to Mali, G. R. et al., (2020).

## Supplemental Movies

**Movie S1**. Coarse grain molecular dynamic simulation of the effect of pulling Dyh3 on ODA remodelling to release shulin.

## Notes

### Competing Interest Statement

The authors have declared no competing interest.

### Summary of Updates

Longer version with more texts and figures. The original version has only 4 main figures. Now, there are 6 main figures and 5 supplementary figures. A lot of text in the supplementary file is moved to the main text. A lot of new text is added as well.

